# Cross-species protection suggests *Entamoeba histolytica* trogocytosis enables complement resistance through the transfer of negative regulators of complement activation

**DOI:** 10.1101/2024.10.04.616735

**Authors:** Maura C. Ruyechan, Wesley Huang, Katherine S. Ralston

## Abstract

*Entamoeba histolytica* is a major cause of diarrheal disease. *E. histolytica* trophozoites (“amoebae”) can invade the intestine and disseminate via the bloodstream, resisting complement lysis through unknown mechanisms. Amoebae kill human cells by performing trogocytosis. After performing trogocytosis, amoebae display human proteins on their own surface and are resistant to lysis by human serum. In this study we sought to further evaluate the mechanism by which amoebae resist complement. To test if complement is responsible for lysis of amoebae, C3-depleted serum was compared to replete serum, and C3 was required for lysis. Amoebae were allowed to perform trogocytosis of human cells and exposed to mouse serum. Although they had performed trogocytosis on a different species than the source of the serum, they were protected from lysis. To test if the protection from lysis by mouse serum was due to the functional interchangeability of human and mouse complement pathway proteins, human CD46 or CD55 (negative regulators of complement activation) were exogenously expressed. Amoebae that expressed human CD46 or CD55 were protected from lysis by mouse serum, indicating that display of human proteins was sufficient to inhibit mouse complement activation. Finally, amoebae were allowed to perform trogocytosis of a cell type in which the complement pathway is not conserved, and they did not become resistant to lysis. Overall, these findings are consistent with the model that trogocytosis enables amoebic acquisition and display of host proteins, including negative regulators of the complement pathway, that provide protection from complement lysis. Since other microbes can perform trogocytosis, this novel mechanism for complement resistance might apply to other infections.

## INTRODUCTION

Amoebiasis is global human diarrheal disease that is caused by the parasite *Entamoeba histolytica* (1). *E. histolytica* cysts are transmitted primarily via fecal contamination of water, and thus, the most affected countries are those with insufficient sanitation (2). Amoebiasis is estimated to cause 67,900 deaths globally per year (3, 4). Because death is a rare outcome of this infection, the global burden of amoebiasis is much higher than the number of deaths per year would suggest. In an endemic area, approximately 80% of infants are infected (5). Malnutrition and stunting are associated with childhood *E. histolytica* infections (6). In children in the Global Enteric Multicenter Study, *E. histolytica* infection was associated with the highest risk of death between enrollment and follow up (7).

In most cases, amoebiasis is asymptomatic or causes mild diarrhea. However, *E. histolytica* can be invasive, causing amoebic dysentery with bloody diarrhea. *E. histolytica* can also spread from the intestine to other organs, resulting in abscess formation in vital organs (2). The liver is most commonly affected, resulting in amoebic liver abscesses that are fatal if not treated. There is no vaccine and therapeutic options are limited (8). There is a limited understanding of the molecular pathogenesis of disease and the determinants of the different disease outcomes.

*E. histolytica* trophozoites (“amoebae”) damage the colon during amoebic dysentery and damage vital organs during disseminated disease, thereby earning their name (*histo-:* tissue, and - *lytic*: destruction or lysis). The profound cell-killing activity of amoebae is likely to be central to tissue damage. Amoebae can kill almost any type of human cell within minutes (9, 10). Amoebic actin is required and amoebae must be intact and viable, since lysates, culture supernatants and killed amoebae all lack cell killing activity (9–11). The previously accepted model was that amoebae secrete pore-forming “amoebapores” (12–15). However, the contact-dependence of cell-killing and the lack of killing activity in cell lysates and supernatants are not consistent with the presence of secreted toxins. Furthermore, transfer of amoebapores to human cells has not been experimentally demonstrated. Cysteine proteases, which are surface-localized and secreted, are key virulence factors that cleave a variety of host substrates, including mucins, antimicrobial peptides, and extracellular matrix components (16–18). Thus, cysteine proteases have a clear role in tissue damage. The surface D-galactose and N-acetyl-D-galactosamine (Gal/GalNAc) lectin is critical for attachment to substrates, including mucins and almost any type of human cell. In disseminated infections, *E. histolytica* survives in the bloodstream, and thus must be capable of evading the host innate immune system to facilitate its passage to the organs. While it is well documented that more virulent strains of *E. histolytica* are better at evading complement lysis, it is not entirely clear what enables this property (19, 20). The heavy chain of the Gal/GalNAc lectin mimics CD59, a negative regulator of complement activation, and is thus one component of complement resistance (21).

In recent years, it has become clear that amoebae kill human cells by performing cell-nibbling, known as trogocytosis (*trogo-:* nibble) (22). During trogocytosis, amoebae pinch off and internalize bites of membrane and intracellular contents from human cells, which eventually results in human cell death (22). Inhibitors and mutants that quantitatively reduce amoebic trogocytosis also reduce human cell death, suggesting that trogocytosis is the mechanism by which amoebae kill human cells (22). In addition to its role in *E. histolytica*, trogocytosis is a nearly ubiquitous form of cell-to-cell interaction in eukaryotic biology, with roles in cell killing, intercellular communication, and cellular remodeling (23). When mammalian immune cells perform trogocytosis, interestingly, proteins acquired from the nibbled cell are displayed by the cell that took bites (24–26).

Through the work of Miller et. al (2019), we found that amoebae that have performed trogocytosis display human cell membrane proteins on their own surface (27). Amoebae that had performed trogocytosis were also protected from lysis by human serum (27). This new finding was consistent with prior work that had shown that amoebae became more resistant to complement after interacting with host cells or tissues, and that this resistance might involve proteins on the amoeba surface. For example, amoebae became more resistant to complement after co-incubation with erythrocytes, and an antibody directed towards an erythrocyte membrane protein reacted with the amoeba surface after co-incubation (28). Animal-passaged amoebae were more resistant to complement lysis than control amoebae(29), and treatment of complement-resistant amoebae with trypsin rendered amoebae complement-sensitive (29, 30).

We hypothesized that protection from lysis by human serum was due to amoebic acquisition and display of negative regulators of the complement pathway. To test this, we created human cell mutants lacking individual complement regulators, but this did not sensitize amoebae to complement lysis, potentially fitting with the redundancy of negative complement regulators (31). When amoebae exogenously expressed individual human complement regulators, CD46 and CD55, this was sufficient for protection from human serum lysis, showing that surface display of complement regulatory proteins can protect amoebae (31). These findings indicated that not only was trogocytosis protective, and it was likely due to the display of human complement regulators (31).

Together, prior studies suggested a model whereby amoebae acquire and display host membrane proteins after performing trogocytosis, and the display of host proteins enables complement resistance (31). However, it was possible that performing trogocytosis generally helps amoebae resist stressors like complement activation and membrane attack complex (MAC) assembly on the amoebic surface. To further test the requirement for display of acquired host proteins in subsequent resistance to complement lysis, amoebae were allowed to perform trogocytosis of human cells and then exposed to mouse serum. Because proteins displayed by amoebae were from a different species than the source of the serum, we did not expect amoebae that had nibbled human cells to resist lysis by mouse serum, but surprisingly, they were protected. Exogenous expression of human CD46 or CD55 protected amoebae from lysis by mouse serum, suggesting that due to the relatedness of these species, aspects of complement regulation are functionally interchangeable. To further test the requirement for display of acquired host proteins in subsequent resistance to complement lysis, amoebae were allowed to perform trogocytosis of a cell type in which the complement pathway is not conserved, and they did not become resistant to lysis by human serum. Together, these findings support that amoebic acquisition and display of host proteins, including negative regulators of the complement pathway, is required for acquired protection from complement lysis.

## RESULTS

### Killing of *E. histolytica* by human serum requires complement activation

We have previously shown that following exposure to human serum, C3b is deposited on the amoeba surface (27). Both C3b deposition and lysis of amoebae are reduced in amoebae that have performed trogocytosis, compared to amoebae that have not performed trogocytosis (27). We also showed that human serum that has been heat-inactivated does not lyse amoebae (27). Together, these findings are consistent with a requirement for complement activation for lysis of amoebae by human serum. To formally test if the complement pathway is required for lysis of amoebae by human serum, amoebae were exposed to C3-depleted human serum or complete human serum, and imaging flow cytometry was used to quantify amoeba viability (Fig.1 and S1). C3-depleted serum led to significantly less amoeba lysis compared to complete human serum (Fig. 1). Thus, lysis of amoebae by human serum requires complement activation.

**Figure 1.** C3 is required for lysis of amoebae by human serum. Amoebae were labeled with CMFDA and exposed to complete human serum, C3-depleted human serum, or control M199s media for 30 minutes. Amoebae were stained with Live/Dead fixable violet and analyzed with imaging flow cytometry. N=6 across 2 independent experiments.

### Amoebae are lysed by mouse serum

We extended our analysis of lysis of amoebae by serum by exposing amoebae to mouse serum. Although mice are not a natural host of *E. histolytica*, experimental infection of mice with amoebae mimics many aspects of the human infection, ranging from immune responses to the host genetic determinants of susceptibility to infection (32). To test if mouse serum led to lysis of amoebae, amoebae were exposed to mouse serum, or human serum as a control, and viability was assessed using imaging flow cytometry. Amoebae were lysed by mouse serum (Fig. S2a). The complement pathway requires Ca^2+^ and Mg^2^ and serum is typically supplemented with Ca^2+^ and Mg^2+^ for *in vitro* studies of complement activation (33, 34). Mouse serum appears to require higher levels of Ca^2+^ and Mg^2+^ supplementation (35), and consistent with this, there was a trend towards higher levels of lysis with higher levels of Ca^2+^ and Mg^2+^ supplementation (Fig. S2a). Mouse serum led to lower levels of lysis overall, in comparison to human serum (Fig. S2b), consistent with its known reduced complement activity (35–36).

### Amoebae that have performed trogocytosis of human cells are protected from lysis by mouse serum

In previous studies, amoebae that had performed trogocytosis of human cells were subsequently protected from lysis by human serum (27). Protection from lysis correlated with display of human proteins on the amoeba surface (31). To further test the requirement for display of acquired host proteins in subsequent resistance to complement lysis, amoebae were allowed to perform trogocytosis of human cells and then exposed to mouse serum. (Fig. 2a-c). We hypothesized that amoebae that had performed trogocytosis of human cells would not be protected from lysis by mouse serum, because the host proteins displayed by amoebae would be from a different species than the source of the serum. However, amoebae that had performed trogocytosis of human cells were protected from lysis by mouse serum (Fig. 2c).

**Figure 2.**
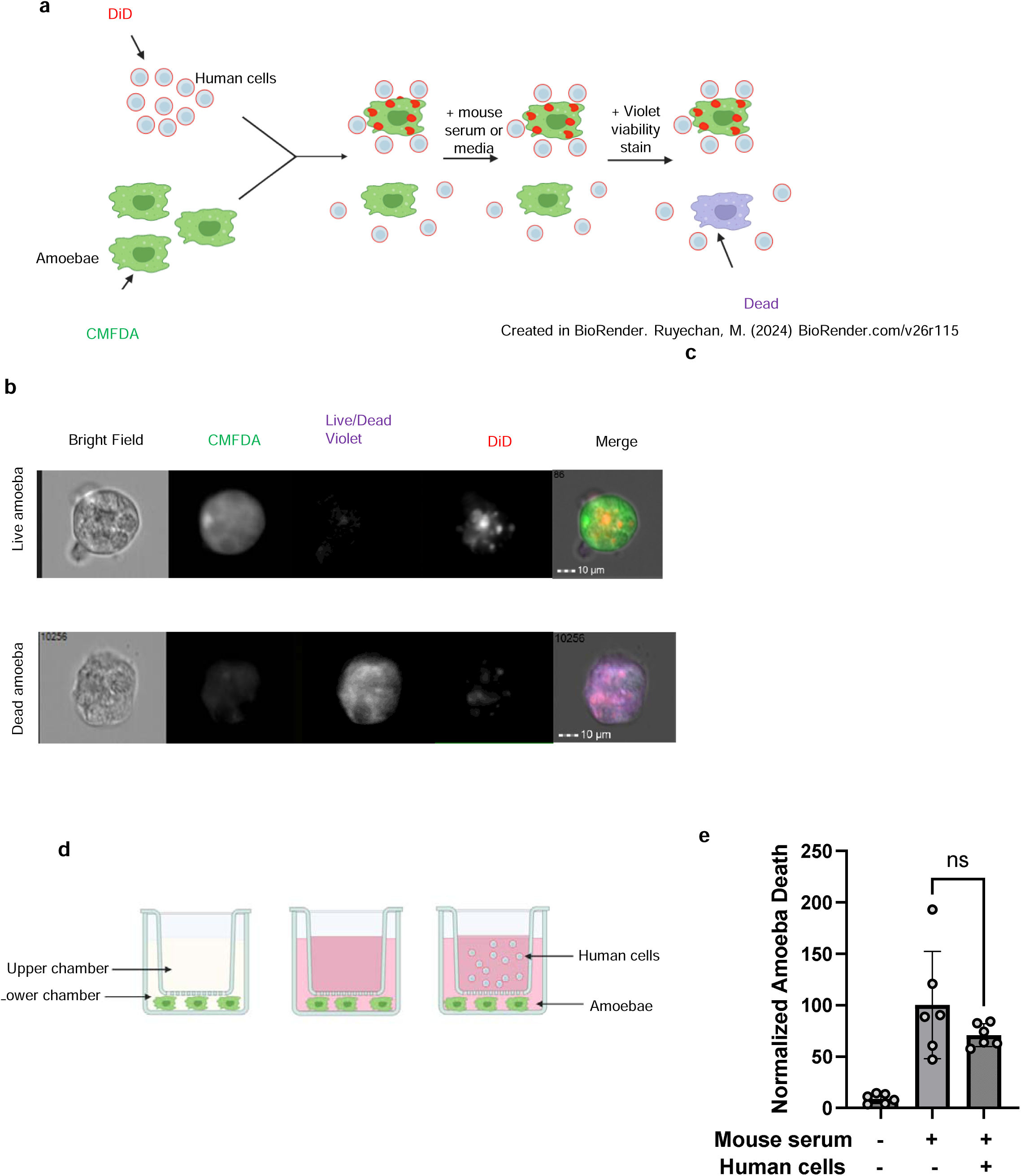
Trogocytosis of human cells protects amoebae from lysis by mouse serum. **a-c,** CMFDA-labeled amoebae were co-incubated with DiD-labeled human Jurkat T cells for 1 hour and then exposed to mouse serum for thirty minutes. Cells were stained with Zombie Violet or Live/Dead fixable violet and analyzed with imaging flow cytometry. N=10 across 3 independent experiments. **b,** Representative images of live and dead amoebae that were exposed to mouse serum following trogocytosis of human Jurkat T cells. **d-e,** A transwell was used to separate amoebae and human cells. CMFDA-labeled amoebae were added to the lower chamber of each transwell, suspended in either control M199s media or mouse serum. To the upper chamber, control media, mouse serum, or DiD-labeled human Jurkat cells resuspended in mouse serum were added. Cells were stained with Zombie Violet or Live/Dead fixable violet and analyzed with imaging flow cytometry. N=5-6 across 2 independent experiments.

The reduced lysis of amoebae by mouse serum could be due to the presence of human cells, which might titrate mouse complement activity. To test if the presence of human cells led to a reduction in mouse serum complement activity, transwell chambers were used. Amoebae were incubated in the absence of human cells in the lower transwell chamber, or human cells were added to the upper transwell chamber (Fig. 2d-e). Trogocytosis is a contact-dependent process, therefore separating the amoebae and human cells into different chambers prevented trogocytosis while still maintaining the same ratio of cells to mouse serum. There was no significant difference in the lysis of amoebae by mouse serum, regardless of whether human cells were present (Fig. 2e). This showed that the presence of human cells does not dilute the complement activity of mouse serum.

### Amoebae that exogenously express human complement regulators are protected from lysis by mouse serum

We initially hypothesized that amoebae that had performed trogocytosis of human cells would not be protected from lysis by mouse serum, because the host proteins displayed by amoebae would be from a different species than the serum. However, because the complement pathway is conserved, it is possible that the displayed human proteins were similar enough to mouse complement proteins to inhibit complement activation. We previously showed that amoebae that exogenously express the negative regulators of complement, CD46 or CD55, are protected from lysis by human serum. Therefore, to ask if human and mouse complement proteins could be functionally interchangeable, we next asked if amoebae exogenously expressing human CD46 or CD55 (Fig. 3a) would be protected from mouse serum. Consistent with our previous findings, exogenous expression of CD46 or CD55 protected amoebae from lysis by human serum (Fig. 3b). Exogenous expression of CD46 or CD55 also protected amoebae from lysis by mouse serum (Fig. 3c-d). This suggests that human and mouse CD46 and CD55 are functionally interchangeable.

**Figure 3.**
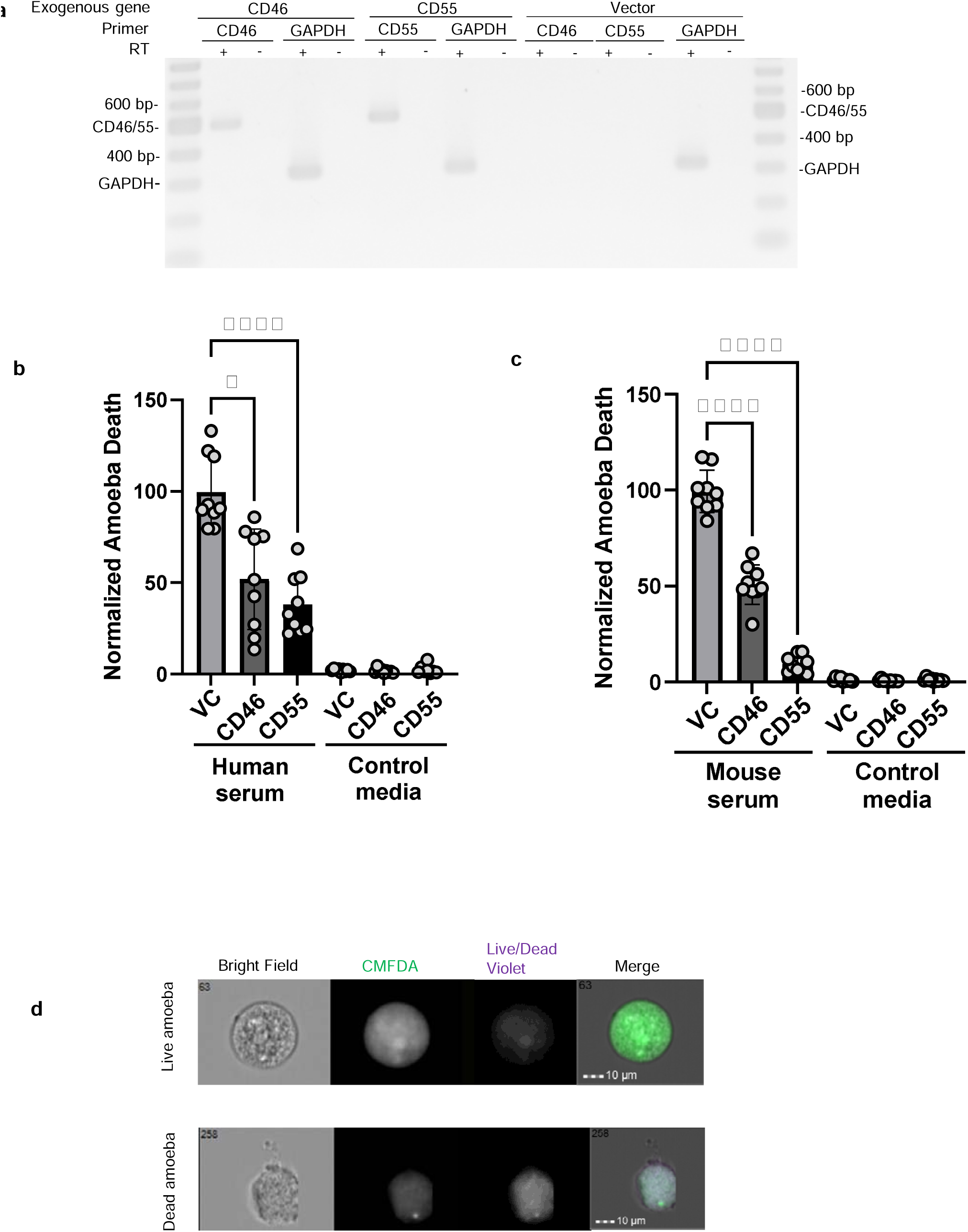
Exogenous expression of human CD46 or CD55 protects amoebae from lysis by both human and mouse serum. **a,** RT-PCR analysis of amoebae stably expressing human CD46, CD55, or the corresponding plasmid vector control. **b-c,** CMFDA-labeled amoebae stably expressing human CD46, CD55, or the corresponding plasmid vector control, were exposed to human or mouse serum for 30 mins. Amoebae were stained with Live/Dead fixable violet and analyzed with imaging flow cytometry. N=8-9 across 3 independent experiments for panel b and N=9 across 3 independent experiments for panel c. **d,** Representative images of live and dead amoebae following incubation with mouse serum.

### Amoebae that have performed trogocytosis of insect cells are not protected from lysis by human serum

To further test the requirement for display of acquired host proteins in subsequent resistance to complement lysis, amoebae were allowed to perform trogocytosis of a cell type in which the complement pathway is not conserved. Consistent with the previously described ability of amoebae to ingest and kill many different cell types (37), amoebae performed trogocytosis of Sf9 insect cells (Fig. 4 and Supplemental Video 1). Amoebae that performed trogocytosis of Sf9 cells were not subsequently protected from lysis by human serum (Fig. 5a). As expected, since the complement pathway is not conserved in Sf9 cells, when Sf9 cells were exposed to human serum as a control, they were efficiently lysed (Fig. 5b). Together, these findings support that amoebic acquisition and display of host proteins, which include functional negative regulators of the complement pathway, is required for subsequent resistance to lysis.

**Figure 4.**
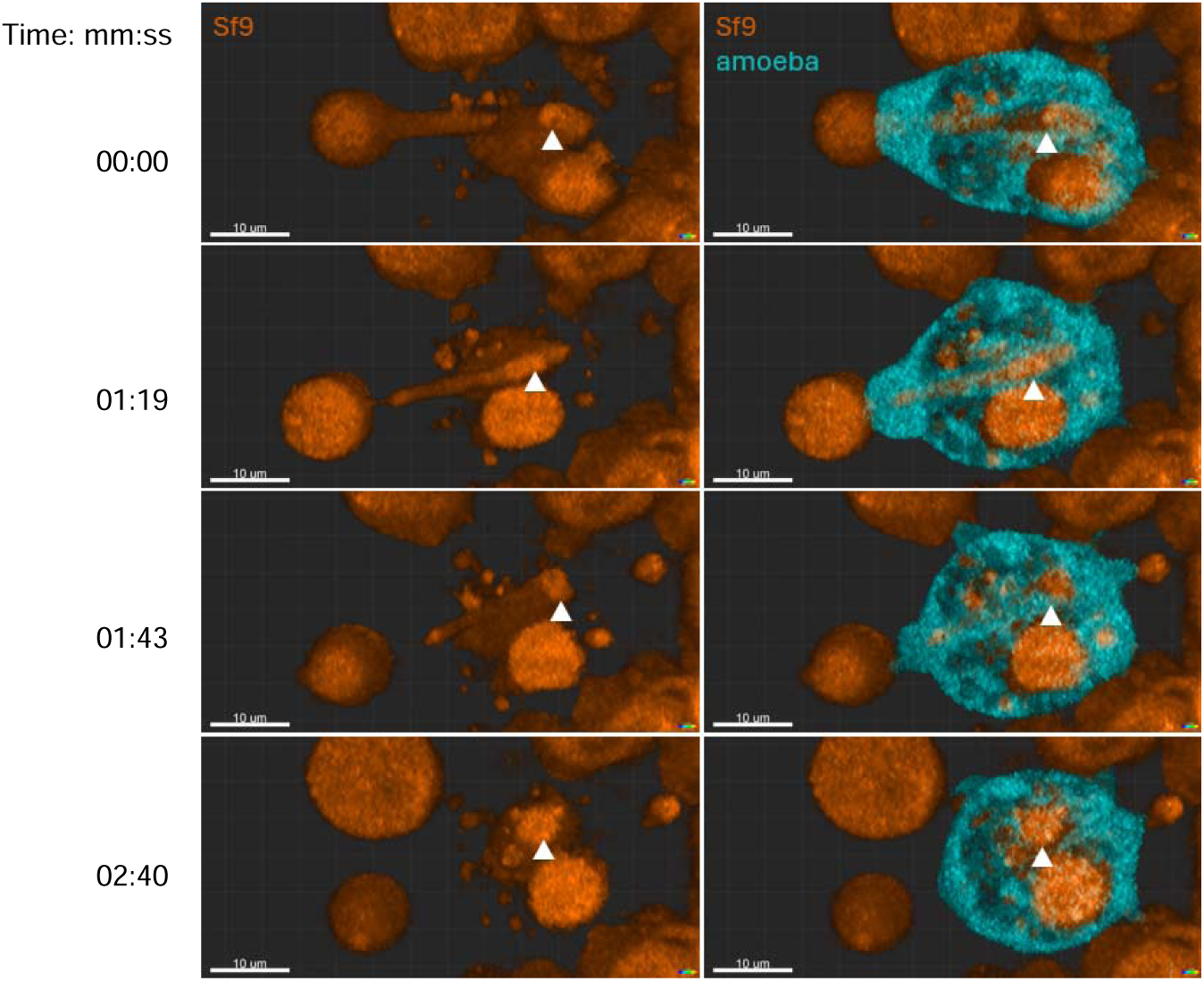
Amoebae perform trogocytosis of Sf9 insect cells. CMFDA-labeled amoebae were co-incubated with CMTPX-labeled Sf9 cells, and imaged live on a Zeiss LSM 980 with Airyscan2. Airyscan functionality was used at the maximum speed to capture rapid z stacks. Images were acquired as a top-down view of entire 3D projection processed and displayed through Blend Rendering Mode in Imaris and Zeiss Zen. The white arrowhead follows a bite of the Sf9 cell as it is nibbled by the amoeba.

**Figure 5.**
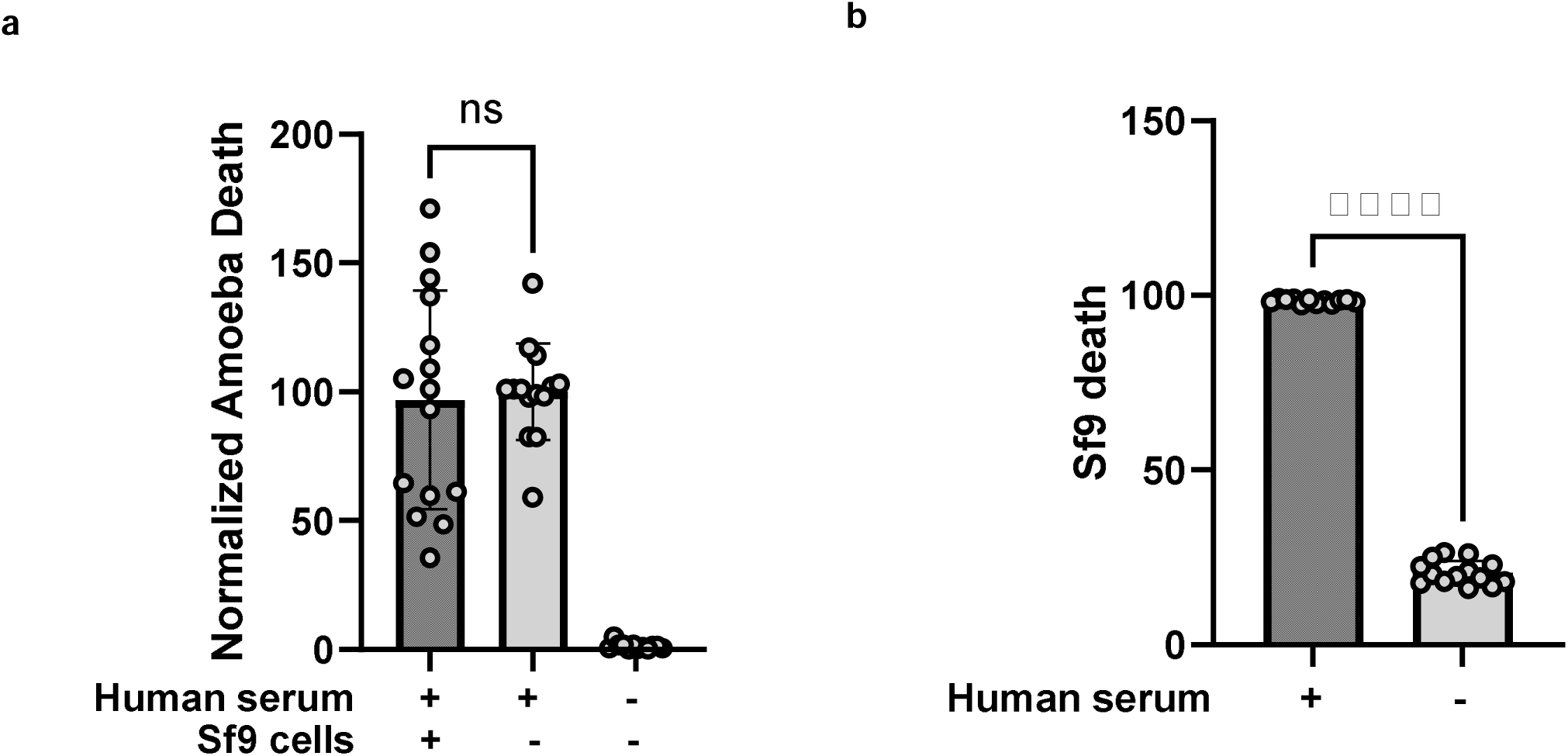
Trogocytosis of Sf9 cells does not protect amoebae from lysis by human serum. **a,** CMFDA-labeled amoebae were co-incubated with CMTPX-labeled Sf9 cells for 1 hour, or incubated in the absence of Sf9 cells, and then exposed to human serum for thirty minutes. Cells were stained with Live/Dead fixable violet and analyzed with imaging flow cytometry. **b,** CMPTX-labeled Sf9 cells were incubated in the absence of amoebae, and then exposed to human serum. N=12-15 across 5 independent experiments.

## DISCUSSION

Prior work suggested that post-trogocytosis display of host cell membrane proteins by amoebae leads to complement resistance (27, 31). To further test this model, in the current studies, amoebae were allowed to perform trogocytosis of human cells and then exposed to mouse serum, and surprisingly, they were protected from complement lysis. We found that amoebae exogenously expressing human complement regulators resisted mouse complement lysis, suggesting that display of human proteins can functionally inhibit mouse complement activation. Amoebae that performed trogocytosis of Sf9 cells, in which the complement pathway is not conserved, did not become resistant to lysis by human serum. Together, these findings support that amoebic acquisition and display of host proteins, including negative regulators of the complement pathway, is required for acquired protection from complement lysis.

In the current studies, C3-depleted human serum did not lyse amoebae (Fig. 1). This is consistent with prior work that showed that heat-inactivation, which inhibits complement proteins, renders human serum unable to lyse amoebae (27). This finding is also consistent with prior studies that showed that amoebae that were killed after serum exposure were intensely decorated with C3b (27). Together, these findings demonstrate that the lysis of amoebae by human serum is due to complement activation.

Here, to further test the requirement for display of acquired host proteins in subsequent resistance to complement lysis, amoebae were allowed to perform trogocytosis of human cells and then exposed to mouse serum. We did not expect amoebae that had nibbled human cells to resist lysis by mouse serum, but surprisingly, they were protected. Exogenous expression of human CD46 or CD55 also protected amoebae from lysis by mouse serum suggesting that due to the relatedness of these species, aspects of complement regulation are functionally interchangeable. Indeed, the mouse and human complement systems are very similar. Both consist of three pathways that act in concert: lectin, classical, and alternative, all converging on the cleavage of C3 into C3b which deposits on the surface of pathogens and sets further immune processes in motion (38). C3b deposition leads to further cleavage of other complement components and the formation of the MAC, that creates pores in the cell membrane and results in cell lysis.

The negative regulators of complement activation are similar in mice and humans. These regulators are critical to prevent the inadvertent activation of the complement system and the lysis of self cells. In both mice and humans, surface-localized negative regulators of complement include CD46 (Membrane Cofactor Protein) and CD55 (Decay Accelerating Factor). CD46 degrades C3b and C4b, while CD55 degrades C3b; thus both prevent the formation of the MAC (39, 40). In mice, CD46 is predominantly found in the testes and retina, and a separate protein, Crry (complement receptor 1-related protein/gene y), is more widely expressed and performs similar functions to both CD46 and CD55 (41, 42). Mouse and human CD46 share ∼48-51% identity and ∼64% similarity, with mouse Crry having ∼34% identity to human CD46 and ∼44% similarity. Mouse and human CD55 protein isoforms share ∼48% identity and ∼47% similarity.

Mouse and human C3 also share ∼77% identity and ∼88% similarity, and mouse and human C4b share ∼76% identity and ∼87% similarity. That the human complement regulatory proteins could provide such protection to amoebae while not having high levels of identity with mouse proteins likely indicates that there is sufficient similarity between these two species’ complement regulatory proteins and their substrates to enable human CD46 or CD55 to inhibit mouse complement activation.

To our knowledge, this is the time that lysis of *E. histolytica* by mouse serum has been evaluated. Mouse serum led to lower levels of amoebic lysis in comparison to human serum (Fig. S2b), These data were consistent with findings from an analysis of the complement activity of wild-type mouse and human serum performed by Latuczek et al (2014) wherein mouse serum had 10 times lower complement activity compared to human serum when tested via a hemolytic CH_50_ assay (36). The reasons for these differences in lytic activity are not currently known, but could potentially be due to differences in the how the different complement pathways function together in each species, as seen in the exposure of dextran-coated SPIO nanoworms to human and mouse serum (43).

Similar to the previous studies in which a transwell was used to separate amoebae and human cells, our findings in Fig. 2e are consistent with a lack of a role for secreted factors in amoebic complement resistance. In our prior work, transwells were used to separate amoebae from human cells, and then amoebae were exposed to human serum (27). Amoebae only became resistant to lysis by human serum if they were incubated on the same side of the transwell as human cells (27). This suggested that secreted factors from human cells, such as proteins or extracellular vesicles, did not have a role in amoebic resistance to complement lysis. Moreover, these findings further affirmed that trogocytosis was likely to be required for complement resistance, since trogocytosis requires direct cell-cell contact. Here, we extended these findings by showing that amoebae that are separated from human cells by a transwell do not resist lysis by mouse serum. This further affirms that secreted factors from human cells are not relevant to amoebic resistance to serum and extends this finding to mouse serum.

Consistent with our previous work (31), amoebae exogenously expressing human CD46 or CD55 were protected from lysis by human serum (Fig. 3b). In our previous work (31), amoebae were stably transfected with plasmids to drive the expression of either CD46 or CD55, and then exposed to increased concentrations of the selective antibiotic (G418) in order to induce overexpression of the exogenous protein. This is a common approach in the *E. histolytica* research community for overexpression (44–47). In the present work, we replaced the promoter in the overexpression construct, with a promoter that drives higher levels of gene expression (48).

This allows for overexpression, without the need to raise the level of the selective antibiotic. This is a preferable approach, since raising the selective antibiotic likely imposes deleterious effects on amoebae. The consistency between our prior findings with overexpression of CD46 and CD55 using raised G418 (31) and the current findings using a stronger promoter (Fig. 3b) demonstrate that the stronger promoter is just as effective. We suggest that this approach is preferable to increased drug selection and is a new tool that is available for the *E. histolytica* research community.

We found that amoebae perform trogocytosis of Sf9 cells. This fits with the known broadness of amoebic cell-killing activity. Sf9 cells are likely to have glycosylated surface proteins that the amoebic Gal/GalNAc lectin could bind, as they have glycosylation pathways which produce n-linked glycoproteins with high mannose glycans and glycans with trimanosyl cores (49, 50). While amoebae were able to attach to Sf9 cells and perform trogocytosis, they did not become resistant to lysis by human serum. The insect immune system does use TEPs or thioester-containing proteins. These function similarly to the mammalian complement system, as complement proteins are a subfamily of thioester-containing proteins, while those in insects are specifically known as insect TEPs (51). Insect TEPs are also noted to be much more similar to α- 2M subfamily of thioester-containing proteins than to complement proteins (51). It is as of yet unknown if there are any insect TEP homologs for complement proteins (51). Our findings indicate that any TEPs present on the Sf9 cell surface are not sufficiently similar to human complement proteins, as there was near total death of Sf9 cells upon exposure to human serum (Fig. 5b).

There are many remaining questions about trogocytosis and immune evasion by *E. histolytica.* While it is evident that host surface proteins are trafficked to the amoebic surface, the pathway by which they are trafficked has not yet been discerned. It is also not known if display of human proteins impacts other aspects of host-parasite interactions, beyond complement evasion. Overall, our findings indicate that trogocytosis by *E. histolytica* enables a novel mechanism for complement resistance, through the display of host negative complement regulators, thereby exploiting the immune system’s own regulatory mechanisms. Since other microbes perform trogocytosis (52, 53), there is the potential for protein display to apply to the pathogenesis of other infections. Further studies of protein trafficking during amoebic trogocytosis may also improve understanding of protein trafficking during eukaryotic trogocytosis in general.

## MATERIALS AND METHODS

### Cell Culture

The HM1:IMSS strain of *E. histolytica* trophozoites from ATCC was cultured as described previously (27, 31, 54). Briefly, amoebae were cultured at 35°C in TYI-S-33 medium supplemented with 80 U/ml penicillin, 80 μg/ml streptomycin (Gibco), 2.3% Diamond vitamin Tween 80 solution (40X; Sigma-Aldrich, or prepared according to Diamond et al, 1978), and 15% heat-inactivated adult bovine serum (Gemini Bio-Products) (55, 56). Amoebae were maintained in either unvented T25 tissue culture flasks or glass tissue culture tubes and passaged when they reached 80% confluence. Amoebae were harvested at 80 to 90% confluence for experiments (27, 31).

Human Jurkat T cells from ATCC (clone E6-1) were cultured as described (27, 31). Briefly, Jurkat cells were cultured at 37°C and 5% CO2 in RPMI 1640 medium (Gibco; RPMI 1640 with L-glutamine and without phenol red) supplemented with 10% heat-inactivated fetal bovine serum (Gibco or Summa Life Sciences), 100 μg/ml streptomycin, 100 U/ml penicillin, and 10 mM HEPES. Jurkat T cells were maintained in T25 vented tissue culture flasks and expanded in T75 vented tissue culture flasks. Jurkat cells were passaged when numbers reached between 5×10^5^ and 2×10^6^ cells/ml, and were harvested at the same cell concentrations for experiments (27, 31).

Sf9 cells from ATCC were cultured at 28°C in Grace’s Insect medium (ThermoFisher) supplemented with 10% heat-inactivated fetal bovine serum (Gibco or Summa Life Sciences). Sf9 cells were maintained in T25 vented tissue culture flasks and expanded in T75 vented tissue culture flasks. Sf9 cells were passaged every 2-3 days at ratios of 1:2 or 1:3.

For the trogocytosis, serum lysis, and live cell imaging experiments (described below), cells were harvested and resuspended in M199medium (Gibco medium M199 with Earle’s salts, L-glutamine, and 2.2 g/liter sodium bicarbonate, without phenol red) supplemented with 5.7 mM L-cysteine, 25 mM HEPES, and 0.5% bovine serum albumin. The supplemented M199 medium is referred to as M199s (27, 31).

### DNA constructs

To make plasmids for exogeneous expression of human CD45 or CD46 in *E. histolytica*, the *E. histolytica* expression plasmid pEhEx (57), was modified to replace the 5’ and 3’ sequences of the cysteine synthase (CS) gene with the 5’ and 3’ sequences of actin, in order to increase the expression level of the exogeneous gene. pEhEx-RLUC, containing a Renilla luciferase (RLUC) insert was used. pEhEx-RLUC was created by PCR amplifying the RLUC sequence from the pcDNA3 RLUC plasmid (58) (Table 1), and the pEhEx plasmid backbone, using Phusion High Fidelity polymerase (ThermoFisher). Gibson assembly with a Gibson Assembly Ultra kit (VWR) was used to ligate the sequences, resulting in the pEhEx-RLUC plasmid.

**Table 1:**
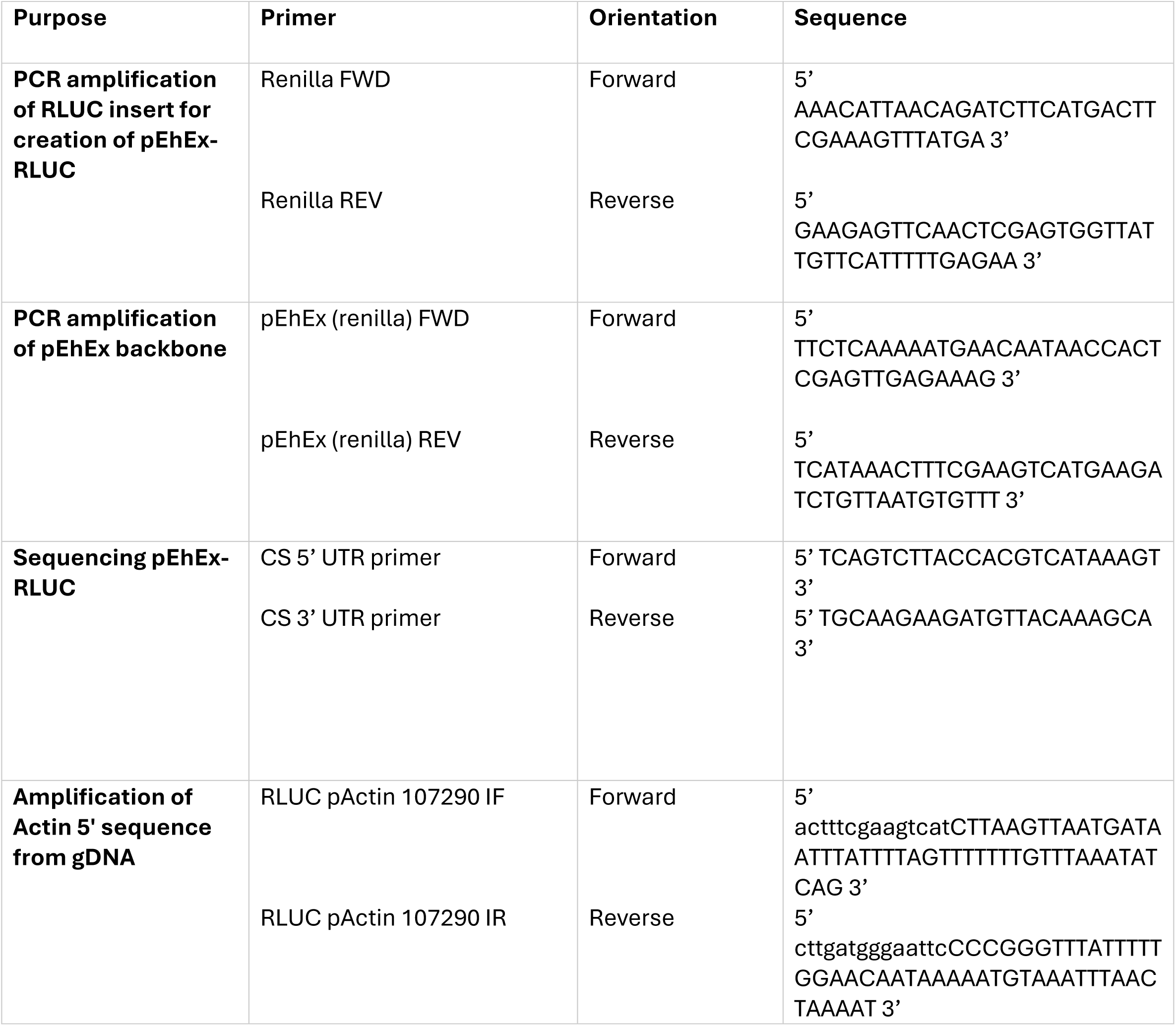
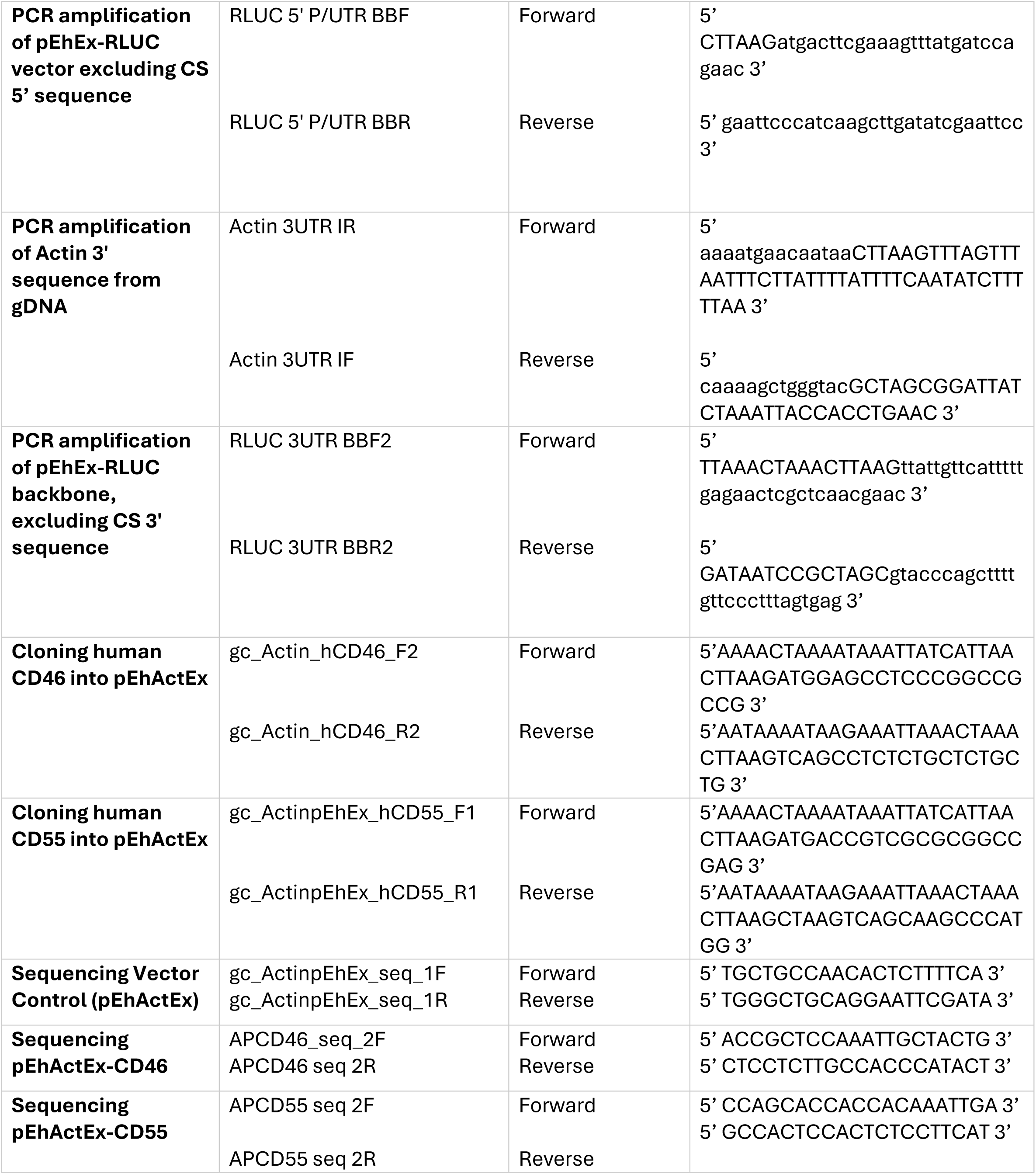
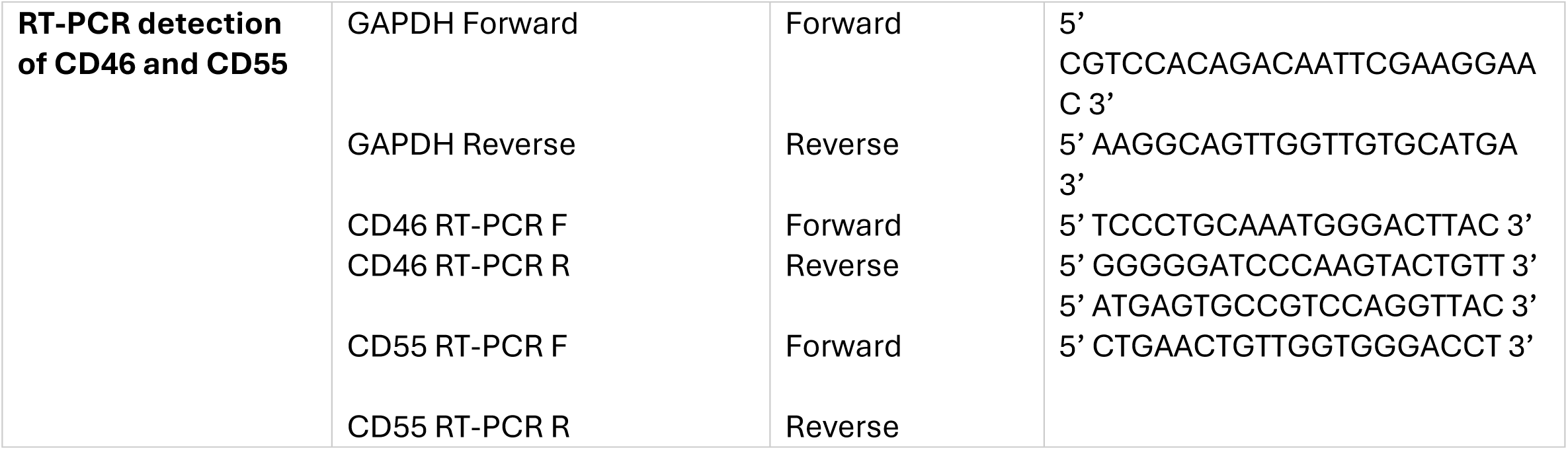
Primers used in these studies.

To replace the CS 5’ sequence with the actin 5’ sequence, the backbone of pEhEx-RLUC was amplified using KAPA high-fidelity polymerase (Roche), while excluding the CS 5’ sequence. A similar process was followed to replace the CS 3’ sequence with the actin 3’ sequence. The actin sequences (upstream and downstream of EHI_107290) were amplified from genomic DNA. Gibson assembly with a Gibson Assembly Ultra kit (VWR) was used to ligate the sequences, creating pEhActEx-RLUC.

Next, the coding sequences for human CD45 and CD55 were PCR amplified from pEhEx-CD45 or pEhEx-CD55 (31). These sequences were originally amplified from cDNA and thus lack introns. The CD46 sequence corresponds to the mRNA transcript variant d (GenBank accession number NM_153826). The CD55 sequence corresponds to the mRNA transcript variant 1 (GenBank accession number NM_000574). pEhActEx was digested with AflII to remove RLUC and the CD46 and CD55 PCR products were each cloned into the backbone using a Gibson Assembly Ultra kit (VWR). The vector control plasmid was created by self-ligating pEhActEx following removal of RLUC by AflII digestion, by using T4 DNA Ligase (NEB). Restriction digest analysis and Sanger sequencing were used to confirm all plasmid sequences. All primer sequences can be found in Table 1.

### Transfection

Amoebae were transfected as described previously (31, 54), by using 20 μg of pEhActEx, pEhActEx-CD46, or pEhActEx-CD55 using Attractene transfection reagent (Qiagen). Stably transfected amoebae were first selected at 3 μg/mL Geneticin (Thermo Fisher Scientific) and then maintained at 6 μg/mL Geneticin (31, 54).

### Gene expression analysis

RNA was extracted using the Direct-zol MiniPrep Plus kit (Zymo Research). RT-PCR was performed as previously described (31), using the primers in Table 1.(31) RNA was extracted once per cell line, and two independent technical replicates of RT-PCR were performed.

### Serum lysis assays

Amoebae were washed in M199s and labeled with CMFDA at 186 ng/ml for 10 minutes at 35°C. Jurkat cells were washed in M199s and labeled with DiD at 21 μg/ml in M199s for 5 minutes at 37°C and 10 minutes at 4°C. CMFDA-labeled amoebae and DiD-labeled Jurkat cells were washed in M199s, combined at a 1:40 ratio in M199s, and co-incubated for 1 h at 35°C. As a control, amoebae were incubated under the same conditions in the absence of Jurkat cells. Next, cells were pelleted at 400x*g* for 8 minutes and were resuspended in 100% C57BL/6 mouse serum (pooled C57BL/6 complement serum; Innovative Research Inc.), 100% human serum (pooled human complement serum; Innovative Research), or M199s depending on the sample and experiment. Mouse serum was supplemented with 75 mM CaCl_2_ and 250 mM MgCl_2_ (35). Human serum was supplemented with 150 μM CaCl_2_ and 500 μM MgCl_2_ (31). Cells were incubated in serum or M199s for 30 minutes at 35°C. Cells were washed and resuspended in M199s and incubated with either Live/Dead fixable violet (Invitrogen) or Zombie Violet (BioLegend) that was prepared as per manufacturer’s instructions at 2 μL/mL for 30 minutes on ice. Samples were fixed with 4% paraformaldehyde for 30 minutes at room temperature. Fixed samples were washed once in 1x PBS, resuspended in 1x PBS and analyzed on an Amnis ImageStreamX Mark II. 10,000 events per sample were collected for samples without Jurkat cells and 100,000 events per sample were collected for samples with Jurkat cells present. Because complement activity in human serum is variable, experiments with at least 3,000 in-focus images of amoebae per replicate (Figure S1d), and average amoeba death in the positive control of at least 10% (Figure S1e) were assessed.

For experiments with Sf9 cells, amoebae were labeled as described above. Sf9 cells were washed in M199s and labeled with CellTracker red CMTPX (Invitrogen) at 590.5 ng/µL for 10 minutes at 28°C. CMFDA-labeled amoebae and CMTPX-labeled Sf9 cells were washed in M199s, combined at a 1:40 ratio in M199s and co-incubated for 1 h at 28°C. As a control, amoebae were incubated under the same conditions in the absence of Sf9 cells. Next, cells were pelleted at 200x*g* for 5 minutes and were resuspended in 100% human serum (pooled human complement serum; Innovative Research) or M199s. Human serum was supplemented as described above.

Cells were incubated for 30 minutes at 35°C in either human serum or M199s. Cells were washed, stained with Live/Dead fixable violet, fixed, washed, and analyzed as described above. 10,000 events per sample were collected for samples without Sf9 cells and 20,000 events per sample were collected for samples with Sf9 cells present. The same cutoffs as above (3,000 in-focus amoebae per replicate and >10% amoeba death in the positive control) were used in the data analysis.

For experiments with amoebae exogenously expressing human proteins, amoebae were labeled with CMFDA as described above. CMFDA-labeled amoebae were washed in M199s and exposed to human or mouse serum supplemented as described above for 30 minutes at 35°C. Cells were washed, stained with Live/Dead fixable violet, fixed, washed, and analyzed as described above. 10,000 events were collected per sample. The same cutoffs as above (3,000 in-focus amoebae per replicate and >10% amoeba death in the positive control) were used in the data analysis.

For experiments with C3-depleted serum, amoebae were labeled with CMFDA, as described above. CMFDA-labeled amoebae were washed in M199s and then resuspended in either M199s, 100% human serum (pooled human complement serum; Innovative Research), or 100% C3-depleted human serum (C3-depleted serum, human; Millipore Sigma) for 30 minutes. Serum was supplemented as described above. Cells were washed, stained with Live/Dead fixable violet, fixed, washed, and analyzed as described above. 10,000 events were collected per sample. The same cutoffs as above (3,000 in-focus amoebae per replicate and >10% amoeba death in the positive control) were used in the data analysis.

### Transwell assays

Amoebae were washed in M199s and labeled with CMFDA at 93 ng/ml in M199s for 10 minutes at 35°C. Jurkat cells were washed in M199s and labeled with DiD at 21 μg/ml in M199s for 5 minutes at 37°C and 10 minutes at 4°C. Mouse serum was supplemented with 75 mM CaCl_2_ and 250 mM MgCl_2_. CMFDA-labeled amoebae were washed in M199s, and resuspended in either mouse serum or M199s. Jurkat cells were resuspended in mouse serum. Amoebae for each condition were placed in the bottom wells of a 12-well transwell plate with 3 μm pores (Corning) and the upper chamber was filled with either M199s, mouse serum, or Jurkat cells resuspended in mouse serum (at a ratio of 1:40 with amoebae). Cells were incubated in a Gaspak bag (BD Biosciences) for 30 minutes at 35°C. Transwells were removed and M199s was added to each well. The plate was then placed back in the Gaspak bag and placed on ice for 5 minutes to detach amoebae for harvesting. Amoebae were washed and resuspended in M199s media and incubated with Live/Dead fixable violet dead cell stain (Invitrogen) or Zombie Violet (Biolegend) that was prepared as per manufacturer’s instructions using 4 μL/mL for 30 minutes on ice. Samples were fixed with 4% paraformaldehyde for 30 minutes at room temperature. Fixed samples were washed once in 1x PBS, resuspended in 1x PBS and analyzed on an Amnis ImageStreamX Mark II. 10,000 events were collected per sample of amoebae exposed to M199s and 20,000 events were collected for samples consisting of amoebae exposed to mouse serum with or without Jurkat cells. Because complement activity in mouse serum is variable, experiments with at least 3,000 in-focus images of amoebae per replicate (Figure S1d), and average amoeba death in the positive control of at least 10% (Figure S1e) were assessed.

### Live Cell Imaging

Amoebae were washed in M199s and labeled with CMFDA at 349 ng/mL in M199s for 15 minutes at 35°C. Sf9 cells were washed in M199s and labeled with CMTPX at 591 ng/mL in M199s for 10 minutes at 28°C. CMFDA-labeled amoebae and CMTPX-labeled Sf9 cells were washed in M199s, and co-incubated in an Attofluor Cell Chamber (Thermo Fisher Scientific) at 35°C without CO_2_ regulation. Cells were initially located using 10-20x objectives and imaged using 63x objectives, on a Zeiss LSM 980 with Airyscan2. Airyscan functionality was used at the maximum speed, capturing cells in the z-plane with an acquisition speed ranging from 80-140 ms. Images were processed with Zeiss Zen imaging software. Processing included a denoise step with a standard length of 1.5-2 to reduce background. The resulting files were converted to Imaris software format and the final video and image processing was performed in Imaris (Oxford Instruments).

### Bioinformatic analyses

Assessment of the relatedness human and mouse complement proteins and negative complement regulators was performed using NCBI BLAST and the available protein sequences from each species. Multiple isotypes from each species were evaluated. The % similarity of these proteins was ascertained using Snapgene again using these same available protein sequences.

### Statistical analyses

Analyses were performed in Prism (GraphPad). Brown-Forsythe and Welch’s ANOVA tests with Dunnett’s T3 multiple comparisons test were used for analyses for Fig. 1, 2c, 2e, 3b, 3c, 5a, S2b and S3, as these assays’ experimental groups did not have equal variances. An unpaired t-test with Welch’s correction was used for analyses for Fig. 5b, as only two groups were being analyzed. *P* values are indicated on each figure: ns, *P* > .05; *, *P* ≤ 0.05; **, *P* ≤ 0.01; ***, *P* ≤ 0.001; ****, *P* ≤ 0.0001.

## ACKNOWLEDGEMENTS

Imaging flow cytometry and microscopy were performed using shared instrumentation in the UC Davis MCB Light Microscopy Imaging Facility. We thank Dr. Michael Paddy and Dr. Thomas Wilkop for technical support in the use of these instruments. We thank the members of our laboratory for helpful discussions. M.C.R. was supported by a NIH National Research Service Award (T32OD011147). This work was supported by NIH R01AI146914 awarded to K.S.R. The funders had no role in study design, data collection and interpretation, or the decision to submit the work for publication. The authors declare no competing financial interests.

## AUTHOR CONTRIBUTIONS

M.C.R. and K.S.R conceived and designed the study. M.C.R. performed experiments and analyzed data. W.H. performed experiments. M.C.R. and K.S.R. interpreted data. M.C.R. and K.S.R. wrote the manuscript.

## SUPPLEMENTARY MATERIAL SUPPLEMENTAL FIGURE LEGENDS

**Supplemental Figure 1.**
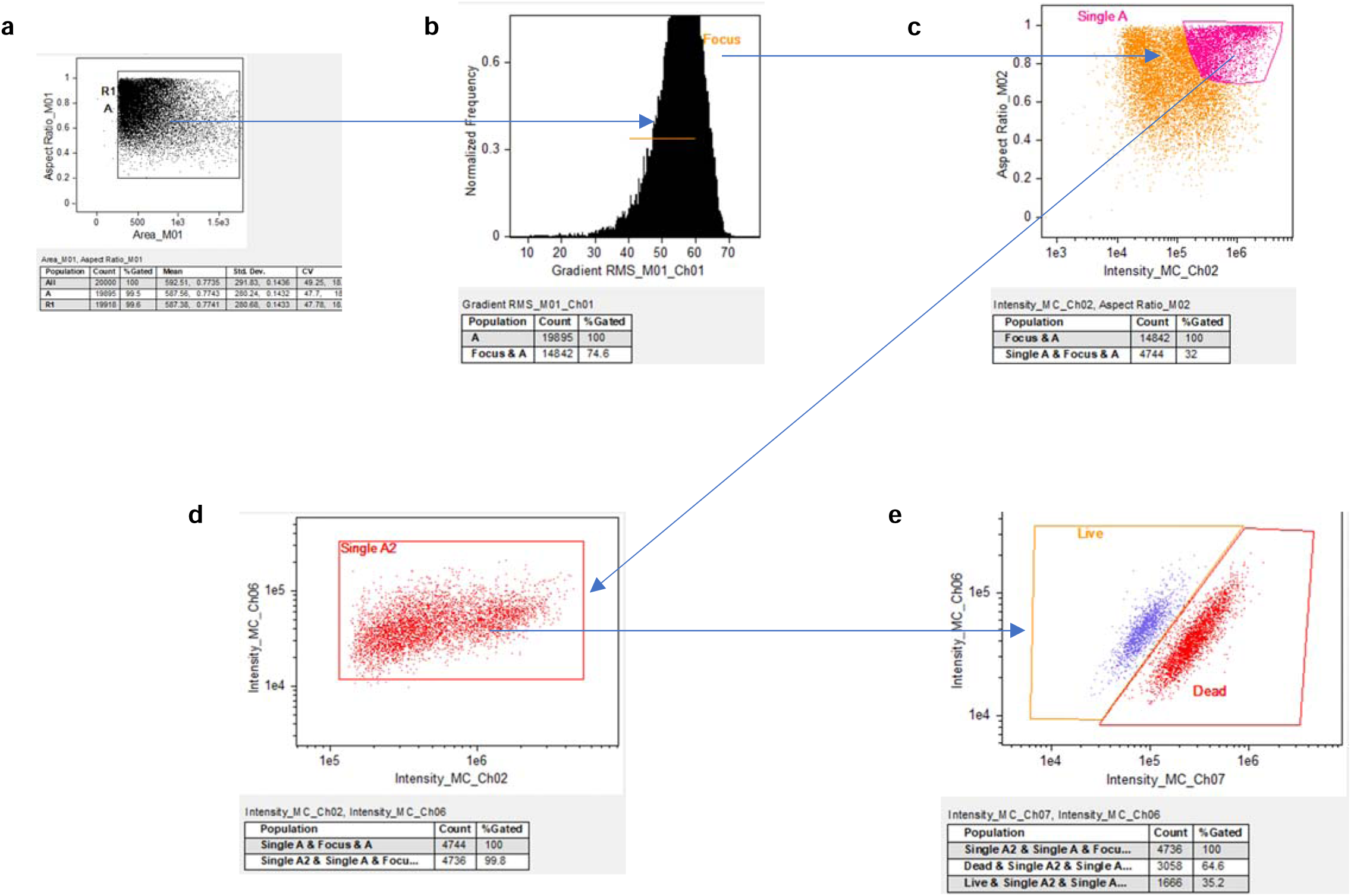
Gating strategy for serum lysis experiments. **a,** All collected images were first gated using aspect ratio and area of the masked brightfield image to remove debris. **b,** Next, images were gated using the gradient RMS of the masked brightfield image to identify images that were in focus. **c,** Images were further gated by aspect ratio and intensity of the channel 2 image (corresponding to CMFDA), and images with single amoebae were gated. **d,** Single amoeba images were further gated by side scatter (channel 6) and channel 2 intensity. **e,** The percentage of amoebae that were dead were then gated by side scatter and channel 7 intensities (Live/Dead violet).

**Supplemental Figure 2. Optimization of mouse serum supplementation with CaCl_2_ and MgCl_2_.** CMFDA-labeled amoebae were exposed to mouse serum for thirty minutes. Amoebae were stained with Live/Dead fixable violet and analyzed with imaging flow cytometry. **a,** To determine the appropriate concentration of CaCl_2_ and MgCl_2_ for mouse serum, amoebae were exposed to multiple different concentrations of both while also using human serum with normal supplementation as a positive control. The maximum amount of CaCl_2_ and MgCl_2_ to be used was based off of the work of Morrison et al (1990). Low corresponds to 15 mM CaCl_2_ and 50 mM MgCl_2_, medium corresponds to 45 mM CaCl_2_ and 150 mM MgCl_2_, and high corresponds to 75 mM CaCl_2_ and 250 mM MgCl_2_. N=2 across 1 experiment. **b,** CMFDA-labeled amoebae were exposed to human serum, mouse serum supplemented with 75 mM CaCl_2_ and 250 mM MgCl_2_, or M199s for 30 minutes. Cells were stained with Zombie Violet dye and analyzed by imaging flow cytometry. N=4 across 1 experiment.

**Supplemental Figure 3. Non-normalized data from Figure 5a. Trogocytosis of Sf9 cells does not protect amoebae from lysis by human serum.** CMFDA-labeled amoebae were co-incubated with CMTPX-labeled Sf9 cells for 1 hour, or incubated in the absence of Sf9 cells, and then exposed to human serum for thirty minutes. Cells were stained with Live/Dead fixable violet and analyzed with imaging flow cytometry. N=12-15 across 5 independent experiments.

## SUPPLEMENTAL VIDEO LEGENDS

**Supplemental Video 1. Live-cell imaging of amoebic trogocytosis of Sf9 cells. CMFDA-labeled**

*E. histolytica* trophozoite performing trogocytosis on CMTPX-labeled Sf9 cells taken using Zeiss LSM 980 with Airyscan2 using multiple z-stacks from a top-down view for 3D imaging.

