## Supplementary figures and images for "Cross-species protection suggests *Entamoeba histolytica* trogocytosis enables complement resistance through the transfer of negative regulators of complement activation"

### Supplemental figures S1-3

#### Slide 1
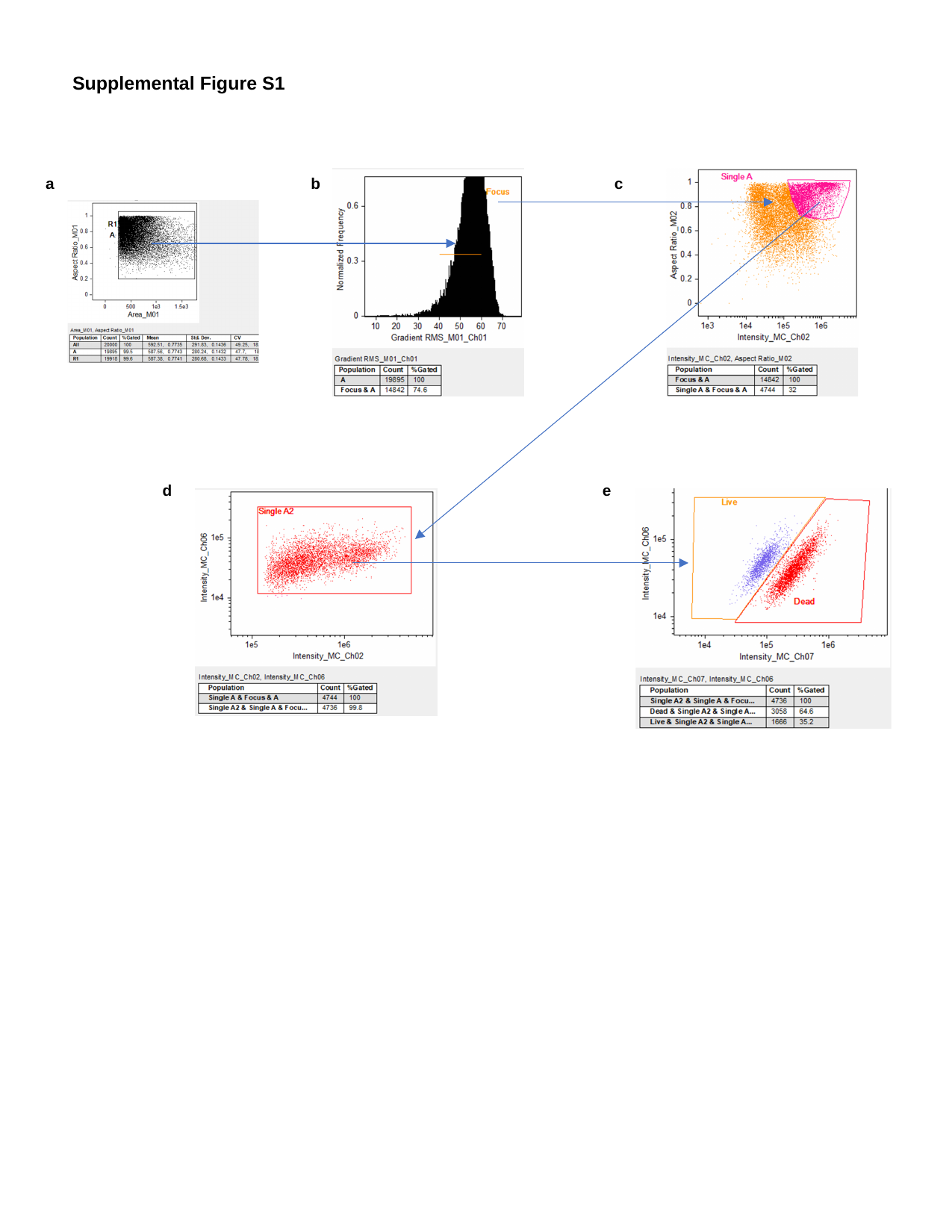

Supplemental Figure S1
a
b
c
d
e

#### Slide 2
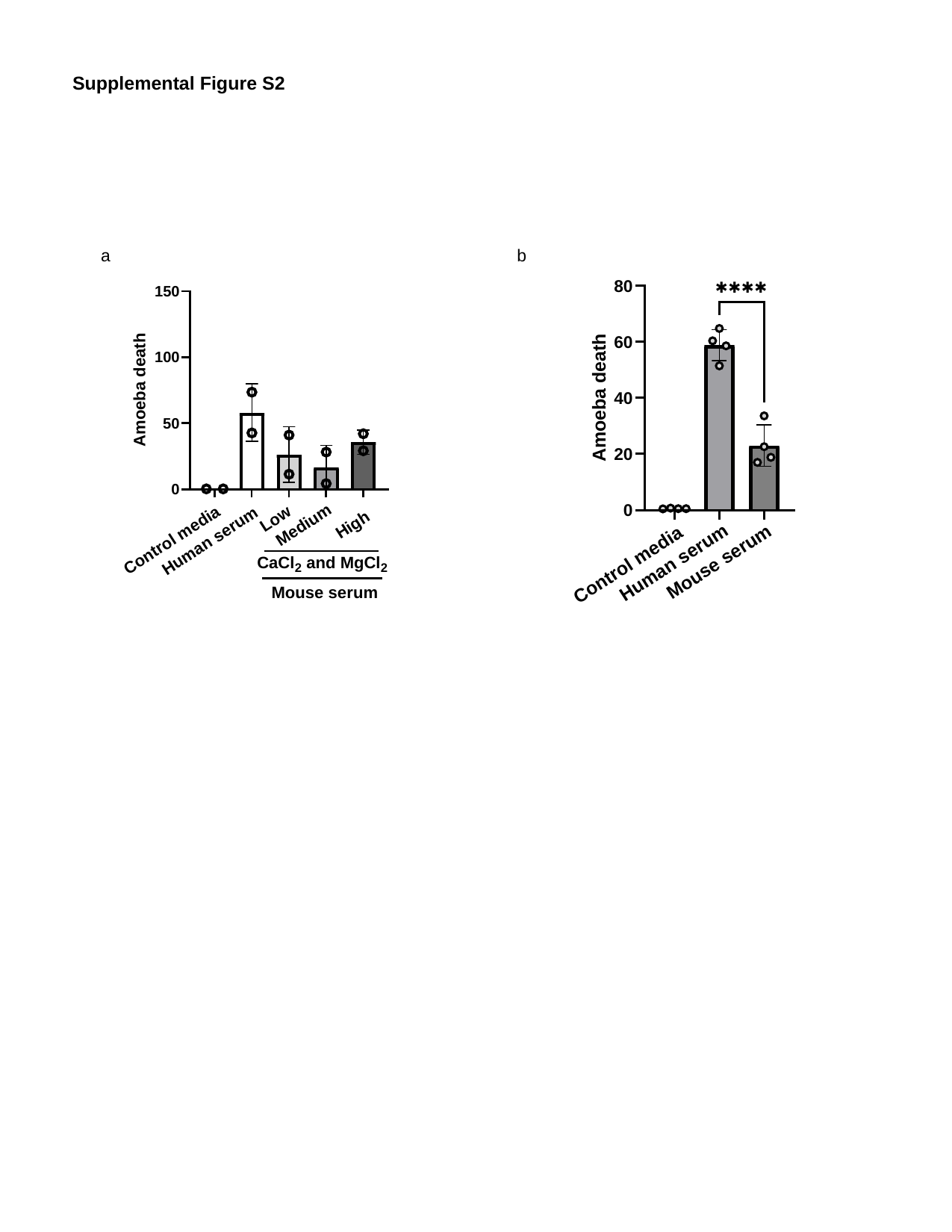

Supplemental Figure S2
a
b

#### Slide 3
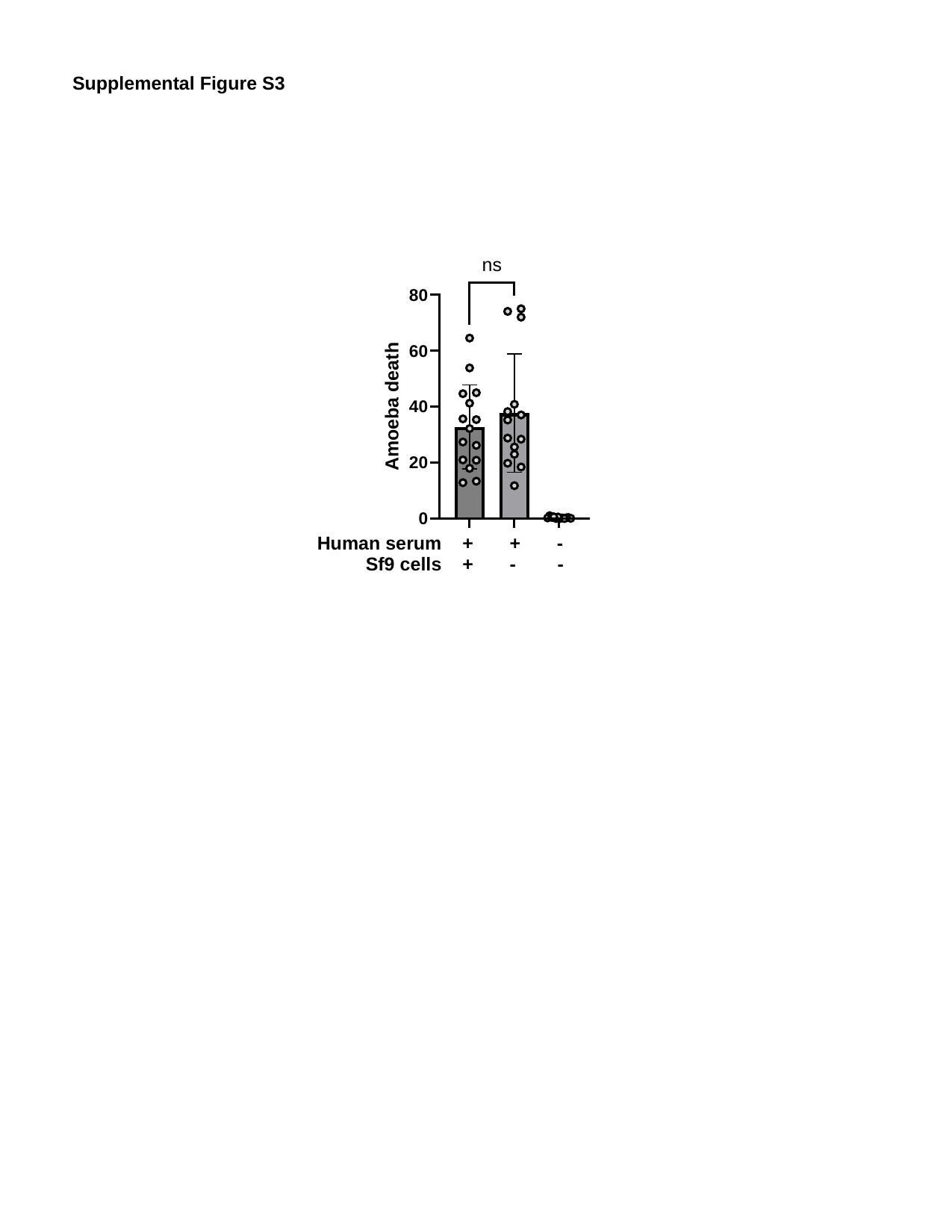

Supplemental Figure S3
